# A dicer-related helicase opposes the age-related pathology from SKN-1 activation in ASI neurons

**DOI:** 10.1101/2023.10.01.560409

**Authors:** Chris D. Turner, Nicole L. Stuhr, Carmen M. Ramos, Bennett T. Van Camp, Sean P. Curran

## Abstract

Coordination of cellular responses to stress are essential for health across the lifespan. The transcription factor SKN-1 is an essential homeostat that mediates survival in stress-inducing environments and cellular dysfunction, but constitutive activation of SKN-1 drives premature aging thus revealing the importance of turning off cytoprotective pathways. Here we identify how SKN-1 activation in two ciliated ASI neurons in *C. elegans* results in an increase in organismal transcriptional capacity that drives pleiotropic outcomes in peripheral tissues. An increase in the expression of established SKN-1 stress response and lipid metabolism gene classes of RNA in the ASI neurons, in addition to the increased expression of several classes of non-coding RNA, define a molecular signature of animals with constitutive SKN-1 activation and diminished healthspan. We reveal neddylation as a novel regulator of the SKN-1 homeostat that mediates SKN-1 abundance within intestinal cells. Moreover, RNAi-independent activity of the dicer-related DExD/H-box helicase, *drh-1*, in the intestine, can oppose the e2ffects of aberrant SKN-1 transcriptional activation and delays age-dependent decline in health. Taken together, our results uncover a cell non-autonomous circuit to maintain organism-level homeostasis in response to excessive SKN-1 transcriptional activity in the sensory nervous system.

**SIGNIFICANCE STATEMENT:** Unlike activation, an understudied fundamental question across biological systems is how to deactivate a pathway, process, or enzyme after it has been turned on. The irony that the activation of a transcription factor that is meant to be protective can diminish health was first documented by us at the organismal level over a decade ago, but it has long been appreciated that chronic activation of the human ortholog of SKN-1, NRF2, could lead to chemo- and radiation resistance in cancer cells. A colloquial analogy to this biological idea is a sink faucet that has an on valve without a mechanism to shut the water off, which will cause the sink to overflow. Here, we define this off valve.

## INTRODUCTION

A central node in the response to xenobiotics and oxidative stress is the cytoprotective transcription factor SKN-1/NRF2, which binds to antioxidant response elements in the promoters of genes, including phase II detoxification and antioxidant synthesis enzymes, that are important for survival under a wide range of stress conditions [1-6]. SKN-1 also plays critical roles in embryonic development by specifying the early blastomere identity and formation of the pharyngeal and intestinal tissues [7-11]. Recently, SKN-1 activity has been demonstrated to coordinate cellular metabolism, particularly lipid metabolism and mobilization [12-18]. Taken together, these studies highlight the complex roles that SKN-1 plays in organismal homeostasis across the lifespan.

Despite the essentiality for development and in the response to stress with age, constitutive activation of SKN-1 has been demonstrated to be pleiotropic for health [13, 15, 17-21]. Although the molecular basis of this health detriment remains elusive, turning off SKN-1 activity is equally important to turning it on. SKN-1 abundance and turnover are regulated by the ubiquitin proteasome system, mediated by the CUL-4/WDR-23 ubiquitin ligase complex. Whereas the loss of *skn-1* is maternal-effect embryonic lethal [7, 22], and whole-life loss of *wdr-23* leads to constitutive SKN-1 activation that results in sickness, the adult-specific inactivation of *wdr-23* can lead to increased lifespan [23]. Relatedly, gain-of-function (gf) mutations in *skn-1* lead to enhanced resistance to acute exposure to oxidative stress early in life, but this stress resistance phenotype is lost when animals reach the post-reproductive period [15]. Tied to these changes in stress resistance and reproduction, these *skn-1gf alleles* also result in the loss of somatic lipid reserves and a significant diminishment of adult lifespan [24] that is tied to the activation of pathogen resistance responses [13]. Considering these discoveries, SKN-1 plays critical roles from embryogenesis through death, which requires sophisticated regulatory mechanisms.

Fifteen years ago, SKN-1 was shown to function in ASI neurons to mediate longevity responses to caloric restriction [25], but multicopy transgenic reporters suggest that SKN-1 activity in non-neuronal tissues is activated in response to stress [11]. Moreover, use of transgenic reporters designed to measure SKN-1 transcriptional responses are activated in cells beyond ASI neurons that suggest that either multicopy transgenics do not accurately reflect physiological SKN-1 responses or that cell non-autonomous pathways responding to SKN-1 activation exists but have yet to be elucidated.

To address these questions, we employed genome editing to tag the endogenous *skn-1* locus with GFP in wild type and *skn-1gf* mutant animals and additionally, replaced the commonly used multicopy extrachromosomal transgenic arrays with single copy tissue-specific constructs that more accurately measure physiological responses. We discovered that SKN-1 expression, even in animals with genetically encoded constitutive SKN-1 activation, remains detectable only in ASI neurons, which is sufficient to coordinate changes in stress and metabolic homeostatic responses. Comparing the transcriptomic landscapes in whole animals, and specifically transcriptional responses to SKN-1 activation specifically in ASI neurons, we further discovered that the pleiotropic outcomes from constitutive SKN-1 activation can be delayed by the dicer-related helicase that can temporarily mitigate the increased transcriptional response from SKN-1 activation.

## RESULTS

### SKN-1 activation phenotypes from two ASI neurons

Previously, activation of SKN-1 has been demonstrated to drive the age-dependent reallocation of somatic lipids such that somatic lipids are depleted while germline lipids are maintained during reproduction that subsequently alters organismal stress resistance and reproduction [15]. In wild type animals, this phenotype is one of the earliest detectable responses upon exposure to pathogenic bacteria like *Pseudomonas aeruginosa* [13]. Although the environment is a potent driver of this response, genetic mutations that activate SKN-1, including gain-of-function (gf) mutations in *skn-1* (Figure S1A) are potent drivers of this change in lipid partitioning [18, 26, 27]; the causality of this single nucleotide mutation on lipid partition was confirmed by genome editing (Figure S1B-C).

To understand how constitutive SKN-1 activation drives premature aging and diminished health, we used CRISPR-Cas9 genomic editing to tag the SKN-1 locus in wild type animals and animals harboring the *skn-1gf(lax188)* allele; hereafter referred to as *skn-1gf*. We used this endogenously tagged version of SKN-1 and SKN-1gf to identify direct transcriptional targets by ChIPseq. RNAseq analyses reveal upregulation of several gene classes including phase II detoxification, host defense factors and lipid metabolism in *skn-1gf* animals. Additionally, significant enrichment of promoters regulating proteostasis factors (e.g., ribosomal subunits, ubiquitin ligases, and chaperones) were recovered by ChIPseq (**Figure 1A-B**). These direct targets included 70 ribosomal subunits (*rps-1* through *rps-27* and *rpl-1* through *rpl-43*), *ubl-1*, *ubq-1*, *lgg-2*, *hsp-1*, *hsp-3*, *hsp-6*, and *ndk-1*, and several genes enriched in neurons (**Figure 1B**).

**Figure 1.**
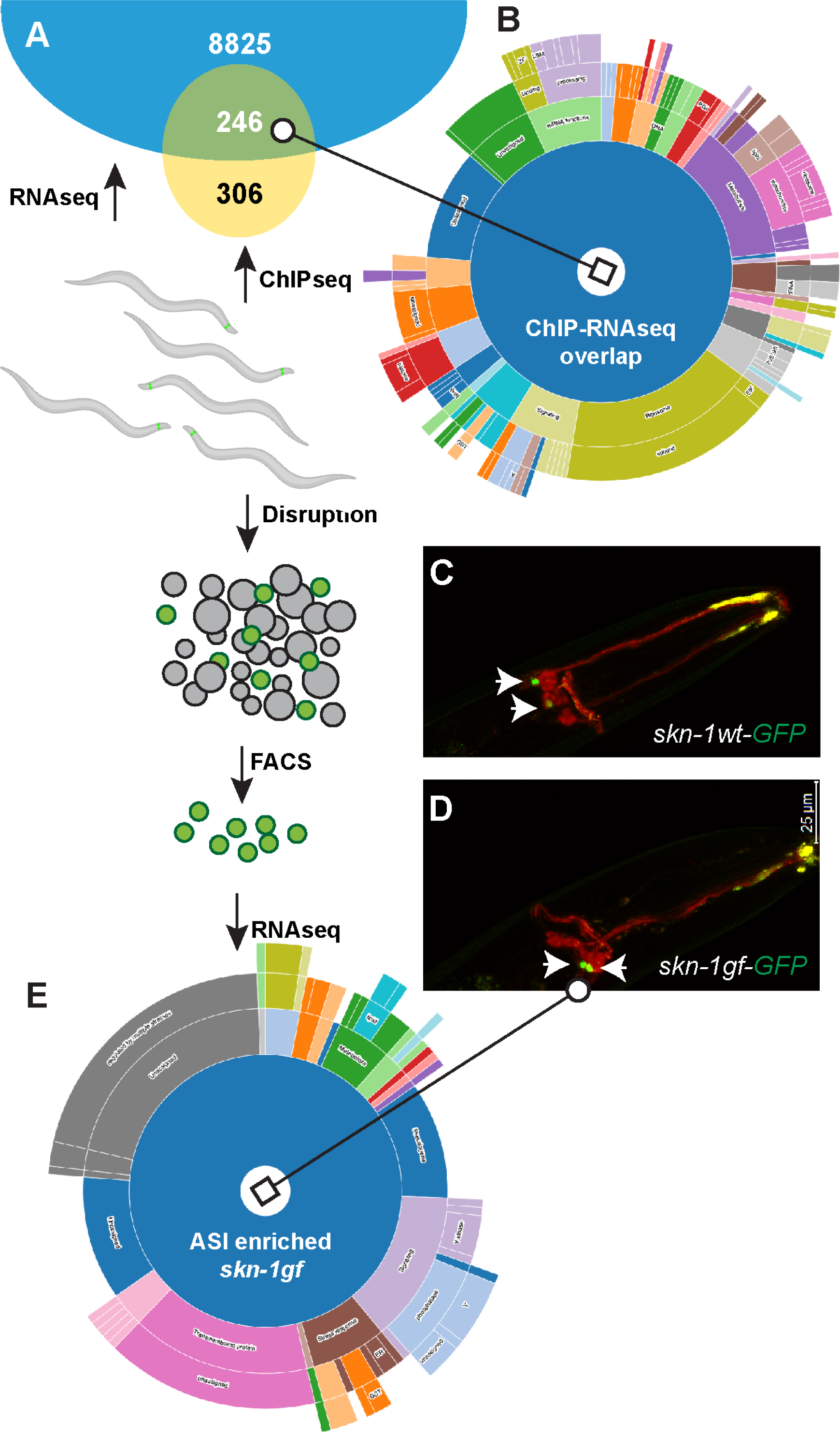
Cell autonomous activity of SKN-1gf in ASI neurons. (**A**) Differentially expressed genes between WT and *skn-1gf* (Blue) were overlapped with SKN-1gf occupied loci (Yellow). ChIP-seq was normalized to a no antibody control IP, while the differential expression analysis was performed by comparing *skn-1gf* to WT transcript levels. Genes with altered expression in *skn-1gf* mutants and identified with occupancy of SKN-1gf on the promoter region were analyzed in WormCat 2.0 (list of genetic loci can be found in Table S1) to reveal enriched classes (**B**). Expression of SKN-1wt-GFP (**C**) and SKN-1gf-GFP (**D**) are indistinguishable and restricted to ASI neurons; co-stained with DiI (red) that marks ciliated neurons; arrows designate GFP in ASI cell bodies (green). (**E**) WormCat 2.0 analysis of genes upregulated in FACS-enriched ASI neuron populations from *skn-1gf* animals.

We investigated whether the low number of recovered targets bound by SKN-1 could be explained by the expression pattern of the SKN-1gf protein. Surprisingly, and unlike animals with activated SKN-1 stemming from loss of the negative regulator *wdr-23* [20, 23, 28-34] or animals exposed to exogenous toxicants [33, 35-37] that have SKN-1 stabilized in the intestine, the expression of SKN-1wt-GFP and SKN-1gf-GFP were only detectable in ASI neurons (**Figure 1C-D**); ASI-specific expression was confirmed by co-expression in animals with ASI neuron expression of mCherry from the *gpa-4* promoter [25, 38](Figure S1N).

Previous transcriptional profiling of animals with activated SKN-1 was performed using RNA samples derived from whole animals. Since SKN-1wt and SKN-1gf protein were only detectable in ASI neurons, we enriched ASI neuronal populations and performed RNAseq. Among the gene targets specifically upregulated in ASI neurons of *skn-1gf* mutants (**Figure 1E**, Table S1) were canonical glutathione s-transferases (*gst-4* and *gst-30)*, pathogen response genes (*irg-5, C55B7.3*), and several neuron enriched genes (*gcy-7*, *C50C3.19*, *nspd-10*, and *zig-2*). Taken together, these data suggest that *skn-1gf* activity may change steady state stress homeostats in ASI neurons.

### Neddylation regulates intestinal SKN-1 stability

SKN-1 coordinates stress adaptation in response to a variety of cellular insults [2, 11-13, 15, 17-20, 39-47] and although *skn-1gf* mutant animals are stress resistant, several details surrounding the molecular basis of this response remain unclear. Traditionally, when a stressor is encountered, SKN-1 protein is stabilized, translocated to the nucleus, and then mediates the transcription of genes that will mitigate the current stress condition [11, 30, 34]. To reconcile the difference in the robust pan-tissue transcriptional response measured in the *skn-1gf* mutant with the inability to detect stabilized SKN-1gf expression outside of the ASI neurons (**Figure 1C-D**), we measured several characteristics of SKN-1 dynamics. First, we tested the stabilization of SKN-1 in response to oxidative stress by acute exposure to hydrogen peroxide (H_2_O_2_), which resulted in the predicted accumulation of SKN-1wt-GFP and SKN-1gf-GFP in the intestine of treated animals (**Figure 2A-B**), while mock treated animals did not display intestinal accumulation (Figure S2N). Despite robust resistance to hydrogen peroxide exposure, we noted a delayed accumulation of SKN-1gf-GFP in the intestine as compared to SKN-1wt-GFP, suggesting an improved capacity to turn over the SKN-1gf protein outside of the nervous system.

**Figure 2.**
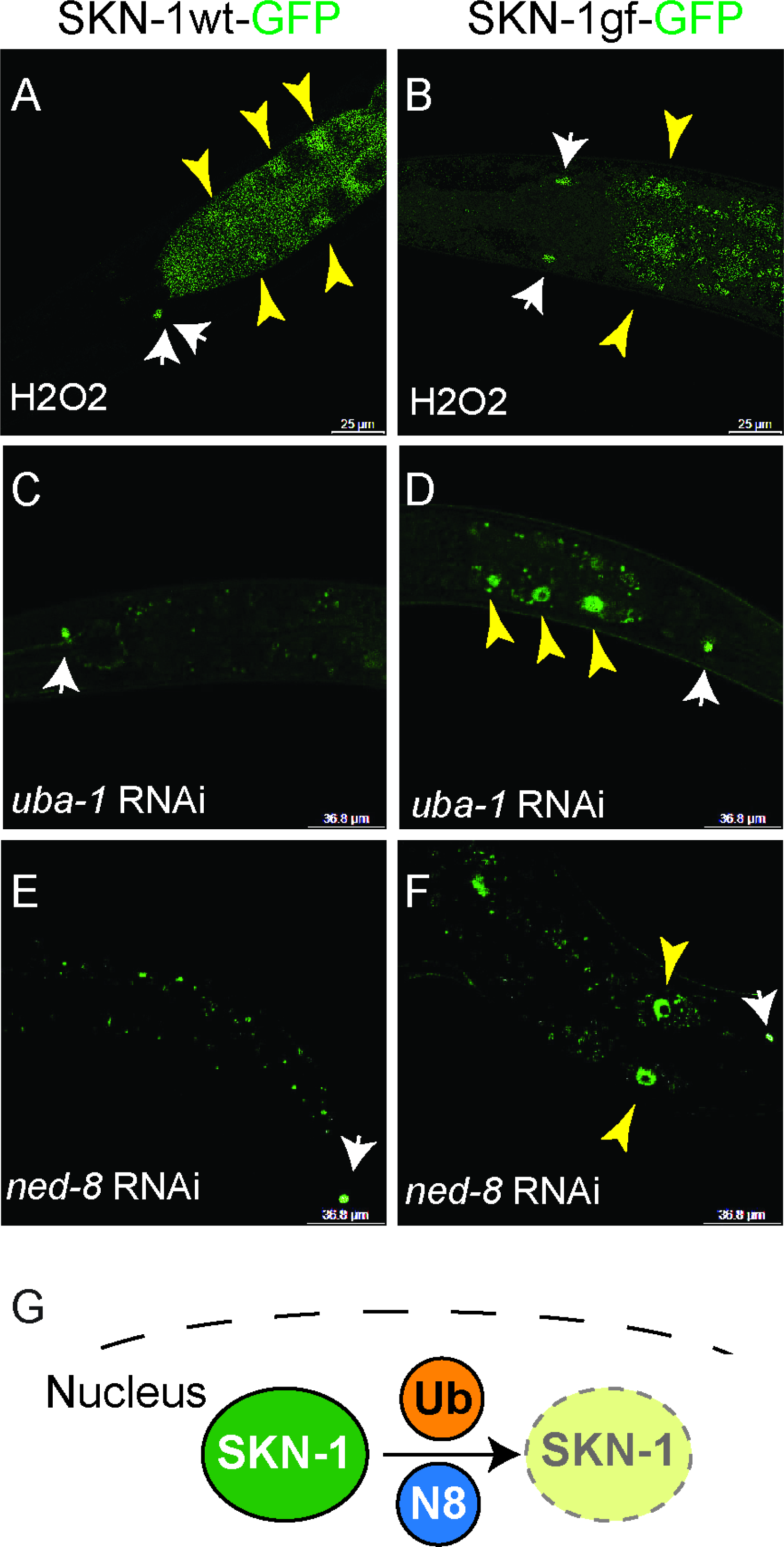
Neddylation regulates nuclear SKN-1 stabilization in the intestine. Acute exposure to hydrogen peroxide drives nuclear accumulation outside of the ASI neurons (white arrows) for SKN-1wt-GFP (**A**) and SKN-1gf-GFP (**B**) within intestinal nuclei (yellow arrows). RNAi of *uba-1* (**C, D**) and *ned-8* (**E, F**) stabilizes SKN-1gf-GFP but not SKN-1wt-GFP in the intestine. (**G**) Cartoon of the role of ubiquitinylation and neddylation on nuclear SKN-1 stability. All RNAi experiments were conducted with n=50 N=3, representative worms are shown.

The stabilization of SKN-1 in response to oxidative stress is linked to its turnover by the CUL4-DDB-1-WDR23 E3 ubiquitin ligase that targets SKN-1 to the ubiquitin proteosome system (UPS) for degradation [20, 34, 42, 48]. As such, we next exposed animals expressing SKN-1wt-GFP or SKN-1gf-GFP to *wdr-23* RNAi and observed accumulation of SKN-1wt-GFP and SKN-1gf-GFP in the intestine (Figure S2A-D). This result confirms that the UPS-mediated control of SKN-1 protein is not perturbed in *skn-1gf* mutants, which supports previous findings that the increase in transcriptional output stemming in *skn-1gf* mutants is additive with loss of *wdr-23* [18].

Although regulation of SKN-1 by the CUL4/DDB1/WDR23 E3 ligase and the ubiquitin proteasome system is well established, the proteostasis network is tightly regulated by the coordinated actions of several pathways [49], including the post-translation modification of proteins by the ubiquitin-like molecules NED-8/NEDD8 [50] and SMO-1/SUMO [51]. We used RNAi to inhibit ubiquitinylation, neddylation, and sumoylation components of the proteostasis machinery to examine their effects on SKN-1wt and SKN-1gf mutants (**Figure 2C-F**, Figure S2E-N). Intriguingly, RNAi targeting *uba-1*, the E1 ubiquitin activating enzyme **(Figure 2C-D**), and *ned-8* (neural precursor cell expressed, developmentally down-regulated 8), the ubiquitin like modifier **(Figure 2E-F**), stabilized SKN-1gf-GFP but not SKN-1wt-GFP within the nucleus of intestinal cells. The preferential stabilization of SKN-1gf-GFP was not observed with RNAi targeting the sumoylation pathway (Figure S2I-N), which suggests the observed differential stabilization of SKN-1gf relative to SKN-1wt is not a generalized sensitivity to proteostatic stress, but instead a specific response to the ubiquitin and neddylation pathways (**Figure 2G**).

### SKN-1gf activity in ASI neurons drives cell non-autonomous decline with age

Our finding that SKN-1gf protein is only detectable outside of the ASI neurons only when proteostasis is perturbed, raised questions as to where SKN-1gf activity is needed to establish early-life stress resistance and eventual health decline if left unchecked. To generate physiologically relevant models for tissue-specific manipulation of SKN-1wt and SKN-1gf activities, we developed several new strains (**Figure 3A**) for tissue-specific expression under established promoter elements as well as tissue-specific degradation by tagging the endogenous *skn-1* locus with an auxin-inducible degron (AID) tag for auxin-mediated degradation of SKN-1wt or SKN-1gf protein exclusively in tissues expressing TIR1 [52].

**Figure 3.**
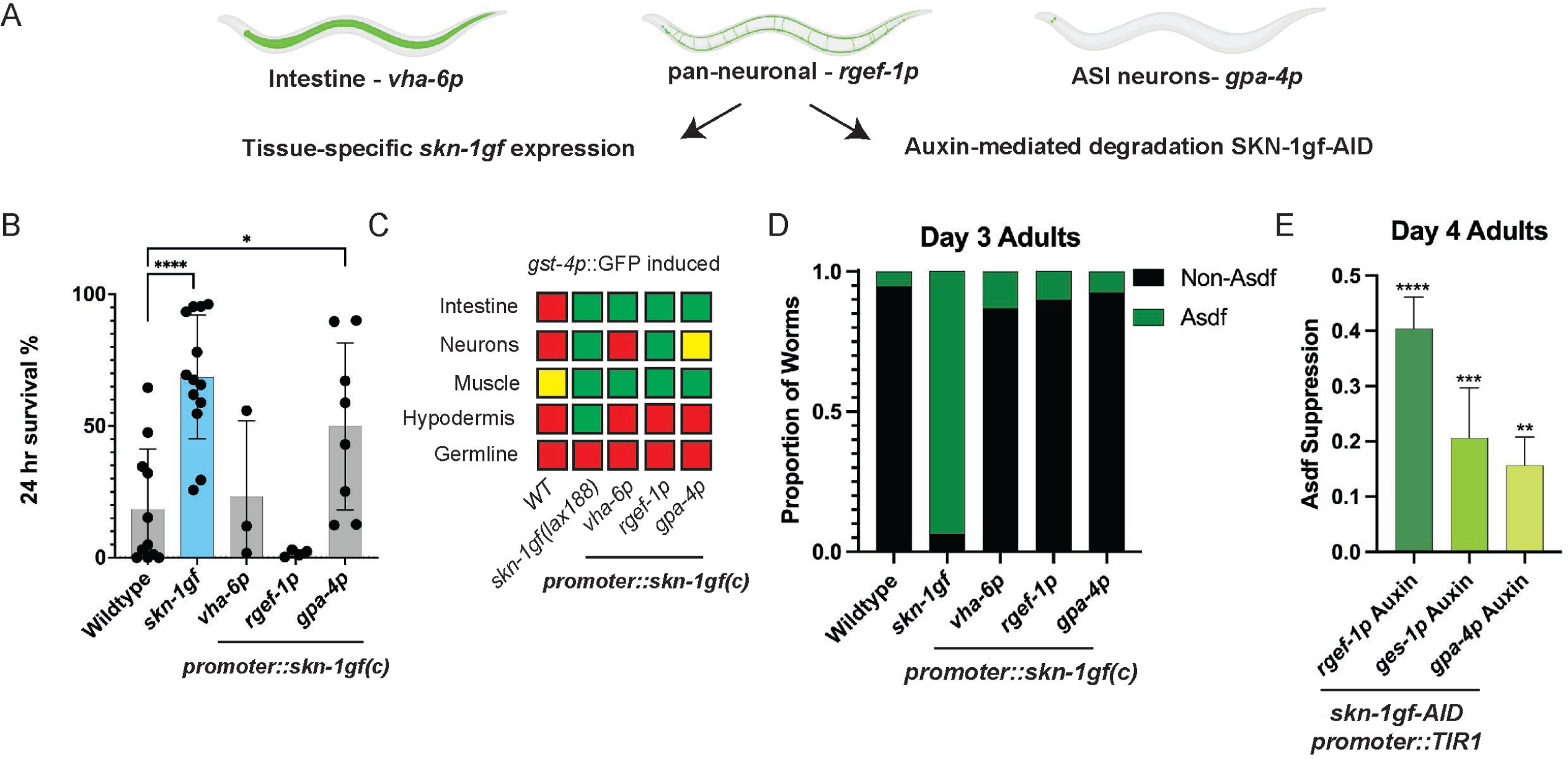
SKN-1 activity in ASI neurons mediates peripheral stress responses. (**A**) Cartoon representation of strains for tissue-specific regulation of *skn-1gf* expression. (**B**) Expression of *skn-1gf* from *gpa-4p* (ASI neurons), but not *vha-6p* (intestine), or *rgef-1p* (pan-neuronal) can establish resistance to acute exposure to H_2_O_2_ n=150 N=3 per condition analyzed by one-way ANOVA *(p<0.05) ****(p<0.0001). (**C**) Tissue-specific expression of *skn-1gf* results in the cell non-autonomous activation of the *gst-4p::gfp* reporter, Green (bright reporter activation), Yellow (dim reporter activation), Red (No detectable reporter activation). Tissue-specific expression of *skn-1gf* is not sufficient to drive age-dependent somatic depletion of fat (Asdf) (**D**). Pan-neuronal, intestinal, and ASI neuron-specific degradation of SKN-1gf can partially suppress somatic lipid depletion (Asdf); n=300; N=3 per condition analyzed by one-way ANOVA **(p<0.01) ***(p<0.001) ****(p<0.0001) **(E)**.

Considering the well-established role that SKN-1 plays in cytoprotection against oxidative stress [11, 53], we tested whether the expression of *skn-1gf* in a single tissue was sufficient to establish resistance to hydrogen peroxide as previously documented for the *skn-1gf* mutant [15]. Both *skn-1B* and *skn-1C* regulate oxidative stress resistance and longevity [11, 25, 54, 55]; however, because the *lax188* gain-of-function mutation does not alter the SKN-1b polypeptide (Figure S1A) we focused on the *skn-1C* isoform. Although previously thought that only SKN-1b activity in ASI was sufficient to drive oxidative stress resistance [11, 25], restricted expression of *skn-1gf* in ASI neurons was sufficient to recapitulate the resistance to acute exposure to hydrogen peroxide (**Figure 3B**), suggesting that the gain-of-function allele is active in ASI and can stimulate organism-level protection from oxidative insult.

The ability of ASI-restricted expression of the *skn-1gf* c-isoform to mediate an organism-level oxidative stress response suggests a cell non-autonomous action stemming from SKN-1 activation. Although we are only able to detect SKN-1wt-GFP and SKN-1gf-GFP in the ASI neurons, our initial isolation of the *skn-1gf* mutant was due to the robust activation of the *gst-4p::gfp* reporter that was induced across multiple tissue types in the animal [18]. As such, we next tested whether tissue-specific expression of *skn-1gf* was sufficient to activate the *gst-4p::gfp* reporter cell non-autonomously (**Figure 3C**, Figure S3A-D). Although muscle-specific expression (*myo-3p*) of *skn-1gf* resulted in muscle restricted activation of *gst-4p::gfp*, ASI specific expression, under the control of the *gpa-4* promoter, induced *gst-4p::gfp* activation beyond the two ASI neurons, including expression in the intestine and body wall muscle. Animals with pan-neuronal expression (*rgef-1p*) displayed a similar pattern of *gst-4p::gfp* activation, except that, multiple neurons displayed activation of the reporter. In contrast, intestine-specific expression (*vha-6p*) resulted in *gst-4p::gfp* reporter activation only in the intestine and body wall muscle and expression of a mCherry reporter alone in ASI was not sufficient to drive the activation of the *gst-4p::gfp* reporter (Figure S3).

We next examined whether cell type specific expression of *skn-1gf* could stimulate the age-dependent somatic depletion of fat (Asdf). Although SKN-1gf activity in ASI neurons was sufficient to recapitulate oxidative stress resistance, it was not sufficient to induce somatic lipid depletion at day 3 of adulthood (**Figure 3D**, Figure S3E-G); however, we did document a trend toward increased somatic lipid depletion in the population at day 5 of adulthood (Figure 3SH-K).

Although we could not identify sufficiency for any single tissue for somatic lipid depletion, we next examined which tissues, if any, were necessary for somatic lipid depletion by auxin-mediated degradation. Based on previous reports with this system [52, 56] we expected and observed some auxin-independent degradation effects on lipid depletion (Figure S3L). However, the addition of auxin greatly enhanced the suppression of lipid depletion in the intestine-specific TIR1 strain by 21%, while pan-neuronal and ASI-specific TIR1 strains resulted in a 40% and 16% suppression of lipid depletion, respectively (**Figure 3E**, Figure S3L-S, Table S2). Collectively, these results reveal that *skn-1* activity in neurons are required for the phenotypes associated with constitutive SKN-1 activation and can initiate a cell non-autonomous response in peripheral tissues like the intestine, which normally removes activated SKN-1 through ubiquitin and neddylation proteostasis pathways.

### DRH-1 activation delays healthspan decline from SKN-1 activity

Our data suggest that activation of *skn-1* in the ASI neurons is sufficient to drive systemic changes in oxidative stress resistance throughout the organism and is needed, at least in part, for the somatic lipid depletion that accompanies SKN-1 activation with age. To identify mediators of the peripheral response to *skn-1gf* activity in ASI neurons, we performed an unbiased genetic screen with ethylmethanesulfonate to recover suppressors of the activation of the *gst-4p::gfp* reporter outside of the nervous system in *skn-1gf* mutants (**Figure 4A**). To our surprise, we recovered a suppressor mutant in the F1 generation of the screen and confirmed the dominant nature of this allele by subsequent backcrossing into the unmutagenized parental strain. We identified insertion-deletion polymorphisms [57] linked to the dominant suppressor mutation that mapped to the center of chromosome IV (LGIV) (**Figure 4B**). Genome-wide sequencing (GWS) of the suppressor mutant genomic DNA revealed twenty-six missense mutations within this region (Figure S4A). Only RNAi targeting the dicer-related helicase gene, *drh-1*, restored the peripheral *gst-4p::gfp* expression observed in the parental *skn-1gf* mutant (**Figure 4C-D**, Figure S4B-G), which also suggests the *drh-1gf(lax257)* mutation is gain-of-function; hereafter referred to as *drh-1gf*. We confirmed causality and dominance of the *drh-1gf* allele by transgenesis (**Figure 4E-F**). The *drh-1* locus encodes for two predicted isoforms, DRH-1A and DRH-1B that are 1037 and 779 amino acids in length, respectively. The *drh-1gf(lax257)* mutation changes glycine 474 in DRH-1A and glycine 216 in DRH-1B to arginine, which are on the surface of the predicted *C. elegans* DRH-1 protein (**Figure 4G**) that resembles the bridging domain found in the mammalian RIG-I that participates in RNA recognition [58, 59].

**Figure 4.**
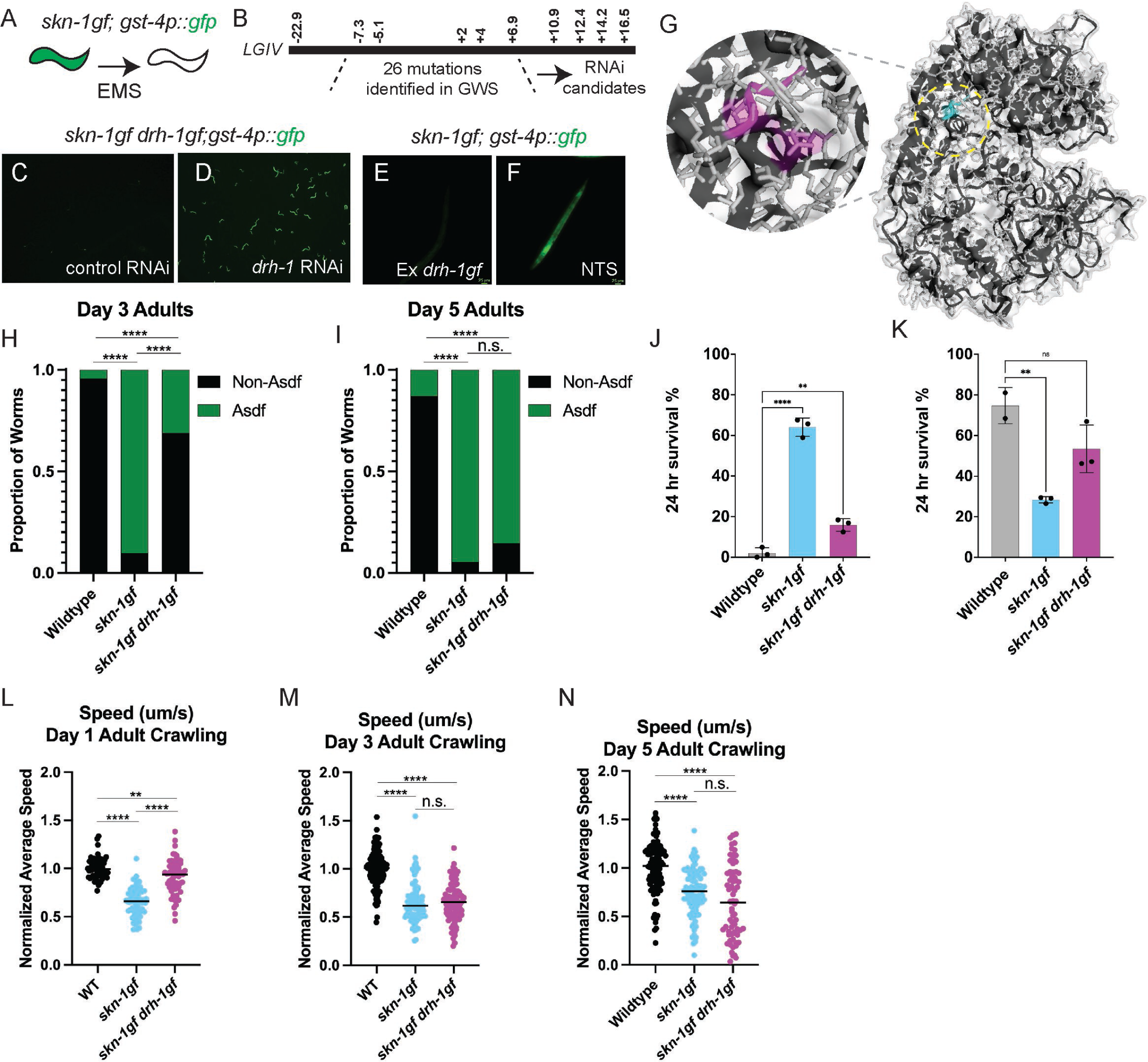
DRH-1 activation delays SKN-1gf-dependent healthspan decline. (**A**) Cartoon schematic of EMS genetic screen for suppressors of *skn-1gf* activation of *gst-4p::gfp*. (**B**) Genetic linkage maps the *lax257* suppressor to LGIV. As compared to control RNAi (**C**) *drh-1* RNAi abolishes the suppression of *drh-1gf* (**D**). Ectopic expression of *drh-1gf* (**E**) suppresses *skn-1gf* activation of *gst-4p::gfp* as compared to non-transgenic siblings (**F**). (**G**) Predicted structure and amino acid substitution (wt-cyan; gf-magenta) in DRH-1gf. *drh-1gf* suppresses the somatic lipid depletion phenotype of *skn-1gf* mutant at day 3 (**H**) but not day 5 (**I**) of adulthood; n=300; N=3 per condition comparisons were made by one-way ANOVA ****(p<0.0001). The resistance to acute H_2_O_2_ by *skn-1gf* exposure at day 1 of adulthood (**J**) and the sensitivity at day 3 of adulthood (**K**) is reversed by *drh-1gf*. (**L-N**) The suppression of the impaired movement phenotype of *skn-1gf* by *drh-1gf* at the L4 larval stage (**L**) is progressively abrogated at day 1 (**M**) and day 3 (**N**) of adulthood; n=50; N=3 per condition. Oxidative stress assay was analyzed by one-way ANOVA **(p<.001) ****(p<.0001), n=100; N=3 per condition.

We next examined the impact of the *drh-1gf* mutation on the age-related healthspan phenotypes influenced by *skn-1gf;* age-dependent somatic lipid depletion (**Figure 4H-I**, Figure S4H-Q, Table S2), oxidative stress resistance (**Figure 4J-K**), lifespan (Figure S4U), and movement (**Figure 4L-N**, Table S3). *skn-1gf drh-1gf* double mutant animals display a significant reduction of somatic lipid depletion at day 3 of adulthood (**Figure 4H**), whereas the phenotype nears complete penetrance in *skn-1gf* mutants. Remarkably, the suppression of somatic lipid depletion was not maintained at day 5 of adulthood where animals harboring the *drh-1gf* allele were indistinguishable from age-matched *skn-1gf* single mutant animals (**Figure 4I**); thus, the effect of the *drh-1gf* mutation is to delay the impact of the *skn-1gf* allele. In addition to changing age-dependent distribution of lipids, the *skn-1gf* mutation has a paradoxical effect on oxidative stress resistance where *skn-1gf* mutant animals are more resistant to acute exposure to oxidative stress at day 1 of adulthood, as compared to WT (**Figure 4J**), but at day 3 of adulthood, when somatic lipid depletion is complete, *skn-1gf* mutant animals are much more sensitive to the same exposure of oxidant than age-matched wild type animals (**Figure 4K**). The *drh-1gf* mutation partially reversed the effects of the *skn-1gf* allele back to wild type (**Figure 4J-K**). The *drh-1gf* mutation did not reverse the shortened lifespan displayed in *skn-1gf* mutants (Figure S4U). Finally, *skn-1gf* mutant animals display movement defects compared to wild type animals that are characterized by reduced speed when crawling starting early in life (**Figure 4L**, Table S3) that persists throughout adulthood (**Figure 4M-N**). Early on, *skn-1gf drh-1gf* mutant animals show an intermediary movement defect that is indicative of partial suppression of the *skn-1gf* mutation which becomes progressively worse and more fully resembles the *skn-1gf* single mutants later in adulthood. Taken together, these results reveal the ability to delay the diminished health stemming from SKN-1 activation with age by an activating mutation in the dicer-related helicase, *drh-1*.

### Intestinal DRH-1 reduces transcriptional response from SKN-1 activation

Universally represented across all genetic mutants with enhanced SKN-1 activity is diminished health with age. Although previous work has demonstrated that the reduced healthspan is associated with the transcriptional activity of SKN-1 [13], our understanding of why constitutive transcription is debilitating requires additional detail. We first examined whether the suppression of the *gst-4p::gfp* transcriptional reporter outside of the nervous system by *drh-1gf* was not an artifact of this simple transgenic reporter. We measured the expression of endogenous *gst-4* transcripts and multiple phase II detoxification genes [60] (**Figure 5A-G**, Table S4), that are strongly induced in *skn-1gf* [13, 18] and regulated by SKN-1 under normal conditions [61, 62] and found that they were significantly reduced in the *skn-1gf drh-1gf* double mutant and their expression pattern opposes the age-related change in the *skn-1gf* single mutant animals.

**Figure 5.**
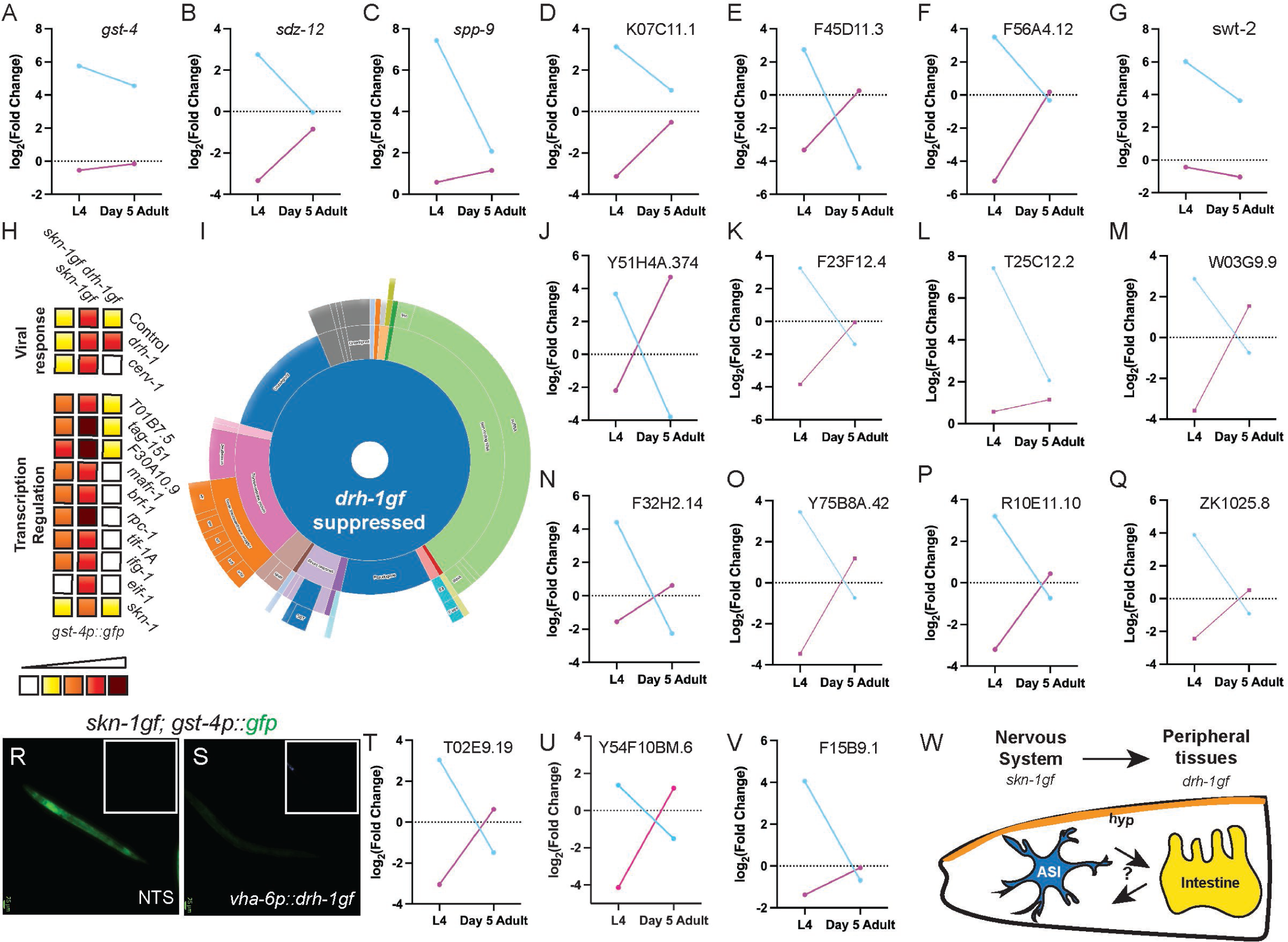
Intestinal DRH-1 activation reduces transcriptional load of activated SKN-1. (**A-G**) *drh-1gf* (magenta) suppresses the activation of established SKN-1 targets in *skn-1gf* mutant animals (blue), blue lines represent differential expression between WT and *skn-1gf* and magenta lines represent differential expression between *skn-1gf drh-1gf* and *skn-1gf*. (**H**) *drh-1gf* mutations abolishes the increased sensitivity of *skn-1gf* mutant animals to RNAi inhibition of transcriptional regulators n=100 N=3. (**I**) WormCat 2.0 analysis of genes activated by *skn-1gf* and suppressed by *drh-1gf*. (**J-Q**) *drh-1gf* suppresses the increased expression of ncRNA in *skn-1gf* mutants. The activation of the *gst-4p::gfp* in *skn-1gf* animals is suppressed by intestinal expression of *drh-1gf* (**S**). (**T-V**) *drh-1gf* suppresses the increased expression of signaling molecules. (**W**) Cartoon schematic of cell non-autonomous signaling by *skn-1gf* in ASI neurons and *drh-1gf* in the intestine. A simple linear regression model was used to compare each of the lines with p<0.05 considered significant.

Next, we examined how *drh-1gf* influences other transcriptional stress responses in the context of constitutive SKN-1gf activity. Previous work identified several RNAi conditions that induce SKN-1 activation [23, 47, 61, 62], including RNAi targeting RNA polymerases themselves and regulators of transcription [61]. RNAi targeting several transcriptional regulators (e.g., *rpc-1*, *F30A10.9*, and *tif-1A)* results in hyperactivation of the *gst-4p::gfp* reporter which does not occur in the context of the *drh-1gf* mutation (**Figure 5H**, Figure S5A). As such, *drh-1gf* attenuates the response to increased transcriptional load resulting from the activation of cytoprotective transcription factors like SKN-1.

DRH-1 is an ortholog of the mammalian retinoic acid-inducible gene I (RIG-I)-like receptors which can detect viral double-stranded RNA (dsRNA) and promote antiviral defense. In *C. elegans*, *drh-1* is required for antiviral RNA interference [63-65], but also plays a separable signaling role in the regulation of transcription immune responses to viral infection [66]. We tested RNAi targeting the CEr1 regulator of viral RNA, *cerv-1*, which strongly induces *gst-4p::gfp* in the *skn-1gf* mutant but not in *skn-1gf drh-1gf* mutants (**Figure 5H**, Figure S5A). Additionally, we confirmed that the reduced activation of the *gst-4p::gfp* reporter was not dependent on the activation of the RNAi-related machinery and as such, unlikely a result of RNAi-induced transgene silencing (Figure S5B). Taken together, these results support a role for *drh-1* in the regulation of SKN-1 dependent transcriptional homeostasis.

In addition to stress response and cellular metabolism genes that are activated in *skn-1gf* mutants but repressed by *drh-1gf* (Figure S5C), we noted an increase in the expression of multiple transcripts enriched for double-stranded secondary structure (e.g., ncRNAs, 21U-RNA, tRNA, snRNA, snoRNA, and pseudogenes) in *skn-1gf* mutants but are repressed in the *skn-1gf drh-1gf* double mutant (**Figure 5I-Q**, Figure S5C). Taken together, these data reveal a new model where activated DRH-1 works to oppose the increased transcriptional load stemming from constitutive SKN-1 activation (**Figure 5W**, Figure S5D). Although the *drh-1gf* allele can temporarily counteract SKN-1 activity, if SKN-1 is not attenuated, the ability of DRH-1 to counterbalance SKN-1 activation is eventually overcome. We noted that the classes of non-coding RNAs were not significantly enriched in the genes upregulated from the ASI neuron population suggesting that the regulation of the ncRNAs occurs outside of the nervous system. As such, we were curious where *skn-1gf* activity was needed to suppress the transcriptional load in the *skn-1gf* mutant background. DRH-1 activity in the intestine has been linked to its RNAi-independent roles in gene expression [66-68]. We expressed *drh-1gf* exclusively in the intestine, which mitigated much of the *skn-1gf*-induced activation of the *gst-4p::gfp* reporter (**Figure 5R-S**). We noted a significant change in the expression of multiple genes that mediate signaling across tissues mediated by the *drh-1gf* allele in the context of the *skn-1gf* background (**Figure 5T-V**) that might mediate the cell non-autonomous responses to activated SKN-1. Our results demonstrate that the genetic activation of the dicer-related helicase opposes the activity of SKN-1 in ASI neurons to reestablish transcriptional homeostasis at the organismal level that improves age-dependent health status (**Figure 5W**, Figure S5D).

## DISCUSSION

The paradoxical discovery that the activation of cytoprotective transcription factors can be the cause of diminished health over the lifespan points to the complexity of the homeostats that have evolved to ensure survival. Our work supports the notion that turning off cytoprotection when it is not needed is perhaps equally as important as our ability to turn it on when it is; analogous to turning a faucet on and off to fill a sink with water. SKN-1 is the *C. elegans* ortholog of the NRF2/NFE2L2 family of transcription factors in mammals. In cancer patients, activation of NRF2/NFE2L2 in transformed cells results in chemo- and radiation therapy resistance [69-71] because the players in phase II detoxification that are normally induced by NRF2 in normal cells to protect against toxic conditions protect the cancer cells from treatment. Beyond this action of the protein products of SKN-1/NRF2, our study revealed that several non-coding RNA transcripts that are induced when SKN-1 is activated may also drive phenotypic decline with age.

Although first reported a decade ago, the molecular basis of how a single amino acid substitution in SKN-1 renders this cytoprotective transcription factor constitutively active remains elusive. Our inability to detect activated SKN-1 stabilized at steady state in cells beyond the ASI neurons was suggestive that either SKN-1 is not functioning in those cell and tissue types or that the *skn-1gf* mutation alters the homeostatic turnover of the gain-of-function protein. Our finding that mildly impairing the ubiquitin proteasome system allows for the detection of intestinal nuclear-localized SKN-1gf but not SKN-1wt supports the latter model. Moreover, the differential hypersensitivity to changes in the ubiquitin-like modifiers of the proteostatic network reveals a previously undescribed role for post-translational modification of neddylation on SKN-1 activity. The stabilization of SKN-1gf in the nucleus when neddylation or ubiquitinylation is impaired supports recent evidence demonstrating that alternating chains of NEDD8-Ubiquitin can mediate nuclear proteotoxic stress [72]. In addition, NEDD8 conjugation to proteins can decrease stability and is an established mechanism to regulate transcription factor function [73] and neddylation can activate cullin ring ligases, like Cul4, through conformational changes which promote cellular proteostasis [74]. Collectively, the observation that proteostatic network has differential effects on SKN-1wt and SKN-1gf is intriguing and suggests the regulation of SKN-1 at the protein level has the capacity for additional regulation.

Beyond the intestine, our work also reveals that while activation of SKN-1 in a single tissue is insufficient to recapitulate all the phenotypes associated with *skn-1gf*, we discovered that neurons are essential for this response. We describe how the activation in the two ASI sensory neurons is required to initiate a cell non-autonomous program in the periphery that drives age-related pathology. Specifically, the activation of SKN-1 in the ASI neurons results in a change in the proteostatic balance within intestinal cells that drives the rapid depletion of intracellular lipid stores in these cells to fuel reproduction [15], which supports the recent model of Morimoto and colleagues that the proteostasis network is critical for reproductive success [75].

While previous work has used transcriptomics to determine differentially expressed genes between WT and *skn-1gf* animals, these results were focused on lipid metabolism and pathogen stress response genes [13]. Here we present a comprehensive multi-omic analysis of animals harboring the *skn-1gf* allele. Previous examinations of SKN-1 in the ASI neuron pair had made observations that it only functioned in determining longevity in a caloric restricted state [25], but our ASI enriched transcriptomic analysis reveals that the repertoire of SKN-1-regulated transcripts includes a larger battery of stress response genes. This finding, in consideration of the apparent lack of SKN-1gf stabilization in other nuclei when additional stressors are absent, further confirms the complexity of the cell non-autonomous nature of the response in ASI neurons and communication needed to peripheral tissues.

Furthermore, our work here provides new insight into the function of SKN-1 isoforms. SKN-1 has four predicted isoforms [11, 54]. SKN-1c has previously been demonstrated to accumulate within intestinal nuclei during oxidative, xenobiotic, and pathogenic stress responses [11, 54, 76] while SKN-1a has a transmembrane domain that allows its association with the mitochondrial [11, 18] and ER [11, 46] membranes that may facilitate organelle specific stress responses. SKN-1b was thought to be the sole isoform accumulating within the ASI neurons and modulating longevity in response to caloric deficit [25], however our findings reveal a gain-of-function mutation that only affects SKN-1a and SKN-1c can drive physiological responses from the ASI neurons and activation of SKN-1a or SKN-1c in the intestine is insufficient. Although the differences in dosage and expression from the single copy-edited genomes used here and multicopy arrays previously used are likely important, taken together, these results suggest that the rigid roles previously assigned to the SKN-1 isoforms are likely more fluid.

We also demonstrate how intestinal activation of the dicer-related helicase, *drh-1*, a regulator of both RNA interference and transcriptional responses to pathogen attack that was previously not connected to SKN-1 activity, can oppose the phenotypes of SKN-1 activation. Animals with constitutive activation of SKN-1 display increased expression of several RNAs with complex secondary structures and are also sensitive to changes in the expression of regulators of multiple RNA polymerases and the cellular proteostasis machinery [47]; a response that is abolished by the *drh-1gf* allele. Thus, DRH-1 acts as a regulator of this RNA homeostat; acting as the drain valve in our analogy and can prevent the sink from overflowing. Intriguingly, activation of DRH-1 only provides a temporary relief to oppose the pathology stemming from SKN-1 activation, since if SKN-1 activity is not turned off, the negative consequences of transcriptional increase still emerge, albeit delayed. Remarkably, the activation of *skn-1gf* in two neurons is sufficient to drive this systemic response that drives multisystem functional decline with age.

Several key questions remain but our results promote the practice of matching exogenous manipulations of cellular cytoprotective responses to physiological signals stemming from cell non-autonomous pleiotropic consequences in distal cells. It remains possible that SKN-1 activation in two or more tissues or that, multiple isoforms, which have different subcellular roles, are necessary to recapitulate the age-related decline in health observed in the *skn-1gf* genetic mutants. The partial mitigation of the *skn-1gf* response when *drh-1gf* is expressed only in the intestine further exemplifies the complexity of SKN-1 signaling and reveals a new layer of cell non-autonomous control in maintaining organismal homeostasis. Moreover, our work provides a framework to refine our models and include alternative approaches that can alleviate the consequences of dysregulated transcriptional activities.

## SUPPLEMENTARY MATERIALS

**Figure S1. Gain-of-function mutations drive SKN-1 activation.** (**A**) Cartoon of genomic location of *skn-1* mutations. A strain with a *de novo* synthesized *skn-1gf* allele but not a strain with the *skn-1gf* allele reverted to WT display somatic lipid depletion at day 3 (**B-D**) and day 5 (**E-G**). SKN-1wt-GFP (**H-J**) and SKN-1gf-GFP (**K-M**) are expressed in ASI neurons (**I, L**) that overlaps with ciliated neurons (**J, M**). *gpa-4p::mCherry* overlaps with SKN-1::GFP in ASI neurons (white arrows) (**N**).

**Figure S2. Ubiquitin-related pathways mediate SKN-1-dependent activities.** (**A-D**) RNAi of *wdr-23* results in nuclear accumulation of SKN-1wt-GFP and SKN-1gf-GFP in the intestine as compared to control RNAi. (**E-N**) RNAi of *ubc-12*, *smo-1*, *uba-2*, *ubc-9* does not result in intestinal accumulation of SKN-1wt-GFP or SKN-1gf-GFP, similar to control RNAi treated animals. Mock treatment of animals with buffer does not induce an oxidative stress response (**M**).

**Figure S3. Cell non-autonomous impact of SKN-1 activation in neurons.** Tissue-specific expression of *skn-1gf* isoform c in the intestine (**A**), muscle (**B**), pan-neuronal (**C**), or ASI neurons (**D**) results in the expression of the *gst-4p::gfp* reporter in multiple tissues. (**E-K**) Tissue-specific expression of *skn-1gf* isoform c does not result in somatic lipid depletion as compared to WT and *skn-1gf* mutant animals (**K**). (**L-S**) Tissue-specific and auxin-dependent degradation of SKN-1gf suppresses somatic lipid depletion (**L**) that does not significantly alter total intracellular lipid levels (**M**).

**Figure S4. Identification and characterization of a *drh-1gf* mutation.** (**A-G**) RNAi screen identifies reducing *drh-1* expression in *lax257* as causal for suppression of *skn-1gf* effects on *gst-4p::gfp* activation. (**H-M**) Representative images of ORO staining revealing *drh-1gf* suppression of day 3 somatic lipid depletion but not day 5. (**N-P**) Representative images of Nile red staining of lipids and quantification at L4 stage; (**Q**). *drh-1gf* mutation does not increase oxidative stress resistance (**R**) and does not increase somatic depletion of lipids (Asdf) (**S-T**). n=100; N=3 per condition; analyzed by one-way ANOVA ****(p<0.0001).

**Figure S5. *drh-1gf* effects are tied to transcriptional regulation.** (**A-B**) *drh-1gf* mutation abolishes the increased sensitivity of *skn-1gf* mutant animals to RNAi inhibition of transcriptional regulators, but the suppression by *drh-1gf* does not depend upon classical regulators of RNAi. (**C**) *drh-1gf* (magenta) suppresses the increased expression of genes induced in *skn-1gf* mutants (blue). (**D**) Cartoon model of the reduction in transcriptional load by *drh-1gf* in the context of unchecked *skn-1gf* activity.

**Table S1**. List of RNAseq genes that overlap with ChIPseq and genes with increased expression in *skn-1gf* ASI neurons. WormCat GO term list with gene list.

**Table S2**. Nile red quantification of lipid abundance.

**Table S3**. WormLab movement data parameters.

**Table S4**. Genes activated by *skn-1gf* that are suppressed by *drh-1gf*.

## MATERIALS AND METHODS

### *C. elegans* Strains and Maintenance

*C. elegans* were raised on 6 cm nematode growth media (NGM) plates supplemented with streptomycin and seeded with OP50. All worm strains were grown at 20°C and unstarved for at least three generations before being used. Strains used in this study include: WT, N2 Bristol strain; SPC168, *dvIs19[gst-4p::gfp*]; *skn-1gf*(*lax188*); SPC572, *dvIs19[gst-4p::gfp*]; *skn-1gf(lax188) drh-1gf(lax257)*; SPC2005, *skn-1*(*lax188syb2619*); SPC2004, *skn-1gf(syb2597)*; SPC2058, *ttTi4348-Pvha-6-skn-1a(lax188)*; SPC2065, *ttTi5605-Prgef-1-skn-1c(lax188)*; SPC2067, *ttTi5605-Pgpa-4-skn-1c(lax188)*; SPC2047, *skn-1gf*-*AID*; DV3803, *ges-1p::TIR1*; DV3805, *rgef-1p::TIR1*; SPC2048, *gpa-4p::TIR1*; SPC597, *Ex[vha-6p::drh-1(lax257)]; skn-1gf; gst-4p::gfp, SPC2094, drh-1gf(syb7642), SPC 600, Ex[gpa-4p::mCherry];skn-1:;GFP, SPC601,Ex[gpa-4p::mCherry];gst-4-4p::gfp*. Single, double, and triple mutants were obtained by standard genetic crosses. Some strains were provided by the CGC, which is funded by NIH Office of Research Infrastructure Programs (P40 OD010440). Genetic mapping of the *drh-1(lax257)* mutation was performed as previously described [18, 19, 39, 40, 42, 77] by linkage of the *gst-4p::gfp* suppression of the *skn-1gf(lax188)* phenotype by using the InDel mapping primer set [57].

### Transgenic and genome editing

CRISPR-Cas9 genome editing was used to revert the *skn-1(lax188)* mutation to WT in *skn-1gf* mutant animals and separately to create an independent gain-of-function allele in wild type animals. To generate tissue-specific expression of *drh-1gf,* the coding and 3’UTR of *drh-1(lax257)* was cloned between the *vha-6p* 5’ regulatory sequence and an SL2::WrmScarlet marker and injected along with a *myo-2p::mCherry* marker into *skn-1gf; gst-4p::gfp* animals. Pharyngeal expression of mCherry was used to identify transgenic and non-transgenic siblings for imaging of the *gst-4p::gfp* reporter.

### RNAi treatment

RNAi treatment was performed as previously described [78]. Briefly, HT115 bacteria containing specific double stranded RNA-expression plasmids were seeded on NGM plates containing 5mM isopropyl-β-D-thiogalactoside and 50μgml^-1^ carbenicillin. RNAi was induced at room temperature for 24 h. Synchronized L1 animals were added to those plates to knockdown indicated genes. RNAi efficiency was determined by qPCR with primers designed for the untranslated region of each mRNA that does not overlap with the dsRNA generated for RNAi; Only RNAi treatments that reduce mRNA levels by 50% were used.

### DiI staining

DiI staining was performed as previously described [79]. In brief, worms were washed with M9 and then put on a rotator overnight in 10ug/mL DiI in M9 at 20°C. Worms were then washed with M9 twice and imaged on an agar pad. All centrifugation was done at 106 rcf for 30 seconds.

### Oil Red O Staining

Oil Red O fat staining was conducted as previously described [57, 80, 81]. In brief, worms were egg prepped and allowed to hatch overnight for a synchronous L1 population. The next day, worms were dropped onto plates seeded with bacteria and raised to 120 h (Day 3 Adult stage) or 168 h (Day 5 Adult stage). Worms were washed off plates with PBS+triton, then rocked for 3 min in 40% isopropyl alcohol before being pelleted and treated with ORO in diH2O for 2 h. Worms were pelleted after 2 h and washed in PBS+triton for 30 min before being imaged at 20x magnification with LAS X software and Leica Thunder Imager flexacam C3 color camera.

For tissue-specific degradation experiments, worms were egg prepped and allowed to hatch overnight for a synchronous L1 population. The next day, worms were dropped onto plates seeded with bacteria and raised to 48 h (L4 stage) and then moved to experiment plates with vehicle 4mM ethanol or 4mM auxin. Worms were moved to new plates every day until 144 h post-drop (Day 4 Adults). Worms were washed off plates with PBS+triton, then rocked for 3 min in 40% isopropyl alcohol before being pelleted and treated with ORO in diH2O for 2 h. Worms were pelleted after 2 h and washed in PBS+triton for 30 min before being imaged at 20x magnification with LAS X software and Leica Thunder Imager flexacam C3 color camera.

### Asdf Quantification

ORO-stained worms were placed on glass slides and a coverslip was placed over the sample. Worms were scored, as previously described [57, 80, 81]. Worms were scored and images were taken with LAS X software and Leica Thunder Imager flexacam C3 color camera. Fat levels of worms were placed into two categories: non-Asdf and Asdf. Non-Asdf worms display no loss of fat and are stained dark red throughout most of the body (somatic and germ cells). Asdf worms had most, if not all, observable somatic fat deposits depleted (germ cells only) or significant fat loss from the somatic tissues with portions of the intestine being clear (somatic < germ).

### Nile Red Staining

Nile Red fat staining was conducted as as previously described [57, 80, 81]. In brief, worms were egg prepped and allowed to hatch overnight for a synchronous L1 population. The next day, worms were dropped onto plates seeded with bacteria and raised to 48 h (L4 stage). Worms were washed off plates with PBS+triton, rocked for 3 min in 40% isopropyl alcohol before being pelleted and treated with Nile Red in 40% isopropyl alcohol for 2 h. Worms were pelleted after 2 h and washed in PBS+triton for 30 min before being imaged at 10X magnification with ZEN Software and Zen Axio Imager with the DIC and GFP filter. Fluorescence is measured via corrected total cell fluorescence (CTCF) via ImageJ and Microsoft Excel. CTCF = Worm Integrated Density-(Area of selected cell X Mean fluorescence of background readings) and normalized to the control.

### Hydrogen peroxide treatment

Conducted as previously described [13]. Briefly, synchronous worm populations at either Day 3 adulthood or 80HPF were washed 3x with M9+.01% Triton. 500uL of 20mM H_2_O_2_ was then added to 600uL of worms+M9+.01% Triton. Worms were then incubated on a rotator at 20°C for 25 minutes before being rinsed 3x with M9+.01% Triton. Worms were then counted and then counted again after 24hrs to determine survival.

### RNAseq Analysis

RNAseq analysis was conducted as outlined [80, 82]. Worms were egg prepped and eggs were allowed to hatch overnight for a synchronous L1 population. The next day, L1s were dropped onto seeded NGM plates and allowed to grow 48 h, 72 h, 120 h or 168 h (L4, Day 1 Adult, Day 3 Adult, or Day 5 Adult, respectively) before collection. Animals were washed 3 times with M9 buffer and frozen in TRI reagent at -80°C until use. Animals were homogenized and RNA extraction was performed via the Zymo Direct-zol RNA Miniprep kit (Cat. #R2052). Qubit™ RNA BR Assay Kit was used to determine RNA concentration. The RNA samples were sequenced and read counts were reported by Novogene. Read counts were then used for differential expression (DE) analysis using the R package DESeq2 created using R version 3.5.2. Statistically significant genes were chosen based on the adjusted p-values that were calculated with the DESeq2 package. Gene Ontology was analyzed using the most recent version of WormCat 2.0 [83]. Simple Linear Regression analysis was done on each time-course slope to determine significant difference between lines, p<0.05.

### Chromatin Immunoprecipitation (ChIP)

Chromatin was prepared as in Nhan *et al.*, 2019 and Wormbook (Modern techniques for the analysis of chromatin and nuclear organization in *C. elegans*). Approximately 1,000,000 L4 synchronized animals were washed in M9 and collected into lysis buffer and flash frozen in liquid nitrogen. Worm pellets were ground via mortar and pestle and resuspended with 1.1% formaldehyde to crosslink proteins. Chromatin was fragmented via sonication and SKN-1::GFP was pulled down by overnight incubation with GFP affinity beads. Associated immunocomplexes were eluted by heat denaturing and DNA fragments were purified. Purified DNA fragments were then used as input for sequencing library preparation using the Diagenode MicroPlex Library prep kit V2. Bioinformatic analyses were done using the following software for ChIP-seq. Quality trimming and Adapter sequences were trimmed from raw paired end reads using Trim Galore package v 0.6.4. Reads were mapped to the *C. elegans* reference genome using bwa v 0.7.17. BAM files were sorted with Samtools v 1.10. BAM files were sorted to contain only uniquely mapped reads using Sambamba v 0.6.8. Peak calls were made using MACS2 v2.2.7.1 Peak files were feature annotated using Chipseeker Bioconductor package in R using the annotate Peak function.

### Statistical Analysis

All statistical analysis was performed using GraphPad Prism version 10.0. Information on specific statistical tests can be found within each figure legend.

### ASI-Enriched RNAseq

Performed as previously described [84, 85]. In brief, approximately 250,000 L4 synchronized WT and *skn-1gf* animals with GFP tagged ASI neurons (*daf-7p::gfp*) were washed with M9 6 times to remove residual bacteria. Animals were then pelleted and Cell Isolation Buffer (20mM HEPES, 0.25% SDS 200mM DTT 3% Sucrose pH8) was added to worms. Worms were incubated in Cell Isolation Buffer for 2min. Initial lysis was quenched by washing with M9 6 times. Worm pellet was resuspended in 20mg/ml Pronase and digested for 20 mins with vigorous pipetting every 5mins through a P1000 tip. Pronase digestion was quenched by resuspending in FBS. Cells were pelleted by centrifuging at 550rcf and resuspended in fresh FBS, Cells were passed through a 40-micron cell strainer. DAPI was added to cells to assess viability. Cells were then sorted on a Bio-Rad S3e FACS system. Neurons were homogenized and RNA extraction was performed via the Zymo Direct-zol RNA Miniprep kit (Cat. #R2052). Qubit™ RNA BR Assay Kit was used to determine RNA concentration. Low input RNA libraries were prepped using the Ovation SoLo RNA library kit from Tecan Genomics. The RNA libraries were sequenced by Novogene. Raw paired-end reads were quality trimmed, and adapter trimmed using timmomatic v0.39. Quality trimmed reads were aligned to the *C. elegans* reference genome using STAR 2.7.6a. Mapped reads were counted via HTseq v2.0.2 union mode. Read counts were then used for differential expression (DE) analysis using the R package DESeq2 created using R version 3.5.2. Statistically significant genes were chosen based on the adjusted p-values that were calculated with the DESeq2 package. Gene Ontology was analyzed using the most recent version of WormCat 2.0 [83].

### Movement Measurements - Crawling

Worms were egg prepped and eggs were allowed to hatch overnight for a synchronous L1 population. The next day, worms were dropped onto plates seeded with OP50. Worms were then allowed to grow until each time point (48 h post-drop for L4s, 72 h post-drop for Day 1 Adults, 120 h post-drop for Day 3 adults, and 168 h post-drop for Day 5 adults). Once worms were at the required stage of development, 30-50 worms were washed off of a plate in 50 uL of M9 with a M9+triton coated P1000 tip and dropped onto an unseeded NGM plate. The M9 was allowed to dissipate, and worms roamed on the unseeded plate for 1 hour before imaging crawling. Crawling was imaged with the MBF Bioscience WormLab microscope and analysis was performed with WormLab version 2022. Worm crawling on the plate was imaged for 1 minute for each condition at 7.5 ms. Worm crawling was analyzed with the software and only worms that moved for at least 90% of the time were included in the analysis. Statistical analysis of crawling speed was done via one way ANOVA with multiple t-test.

### Protein prediction

Phyre2 [86] was used to predict the structure of DRH-1wt and DRH-1gf; WT structure prediction file 25950237 and Missence3D (image with cyan and magenta in same structure, predicted changes in protein structure 30995449).

## ACKNOWLEDGMENTS

We thank J. Gonzalez, M. Lynn, and S. Ledgerwood for technical assistance. This work was funded by the NIH R01AG058610 and Hevolution Foundation award HF AGE-004 to SPC, F31GM137587 to CDT, F31AG077873 to NLS, and T32AG052374 to NLS and BTVC. We also thank the USC School of Gerontology Imaging Core that is funded in part by the Nathan Shock Center of Excellence P30AG068345. Some strains were provided by the CGC, which is funded by the NIH Office of Research Infrastructure Programs (P40 OD010440). We thank WormBase for database curation and data access.

## Author contributions

Conceptualization: SPC; Methodology: CDT, NLS, CMR, BTVC, and SPC; Investigation: CDT, NLS, CMR, BTVC, SPC; Visualization: CDT, NLS, CMR, BTVC, SPC; Supervision: SPC; Writing (original draft): CDT and SPC; Writing (reviewing & editing): CDT, NLS, CMR, BTVC, SPC.

## Data and materials availability

All data are available in the main text or the supplementary materials.

