## Supplementary figures and images for "A dicer-related helicase opposes the age-related pathology from SKN-1 activation in ASI neurons"

### Supplemental Figures

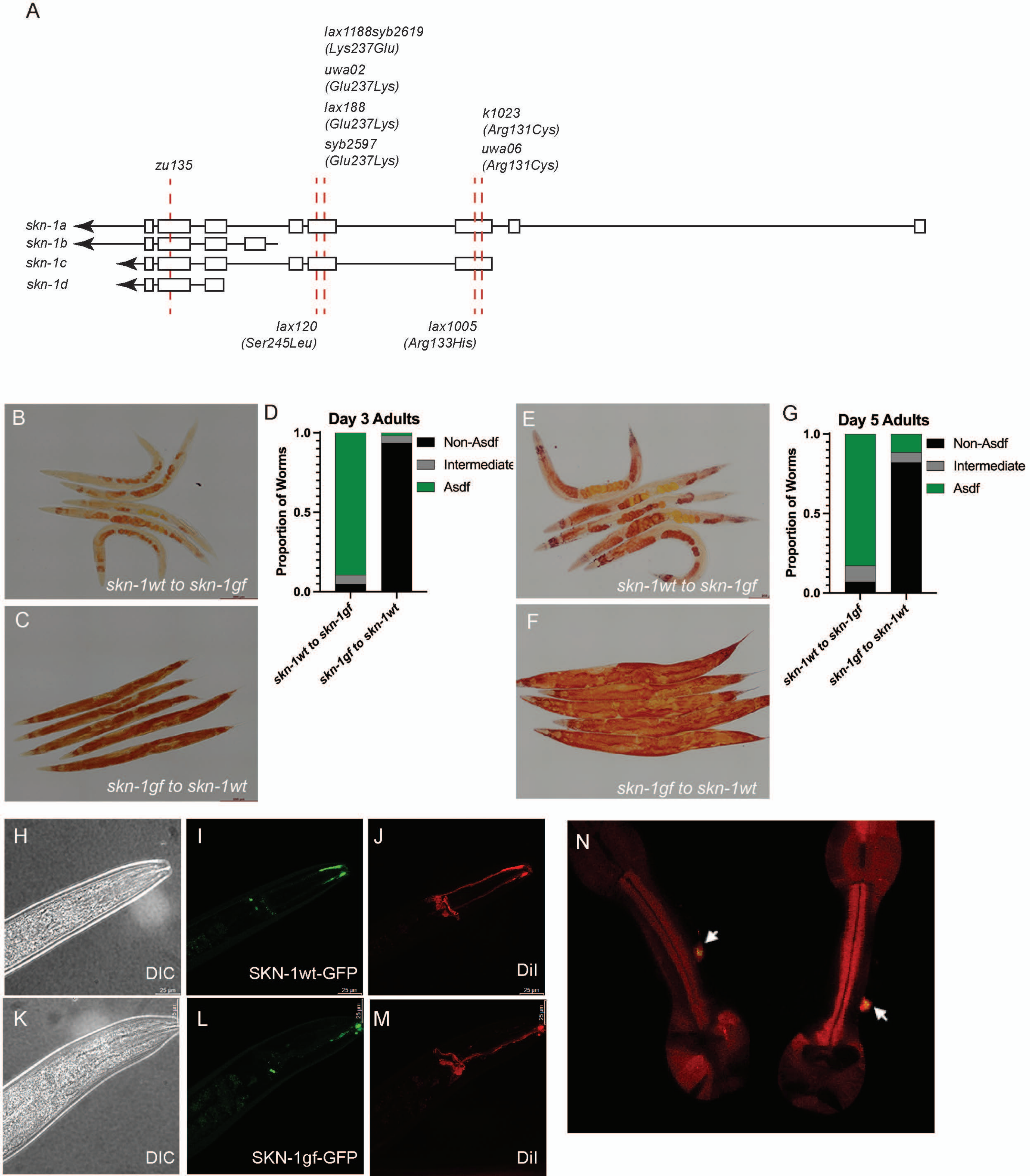

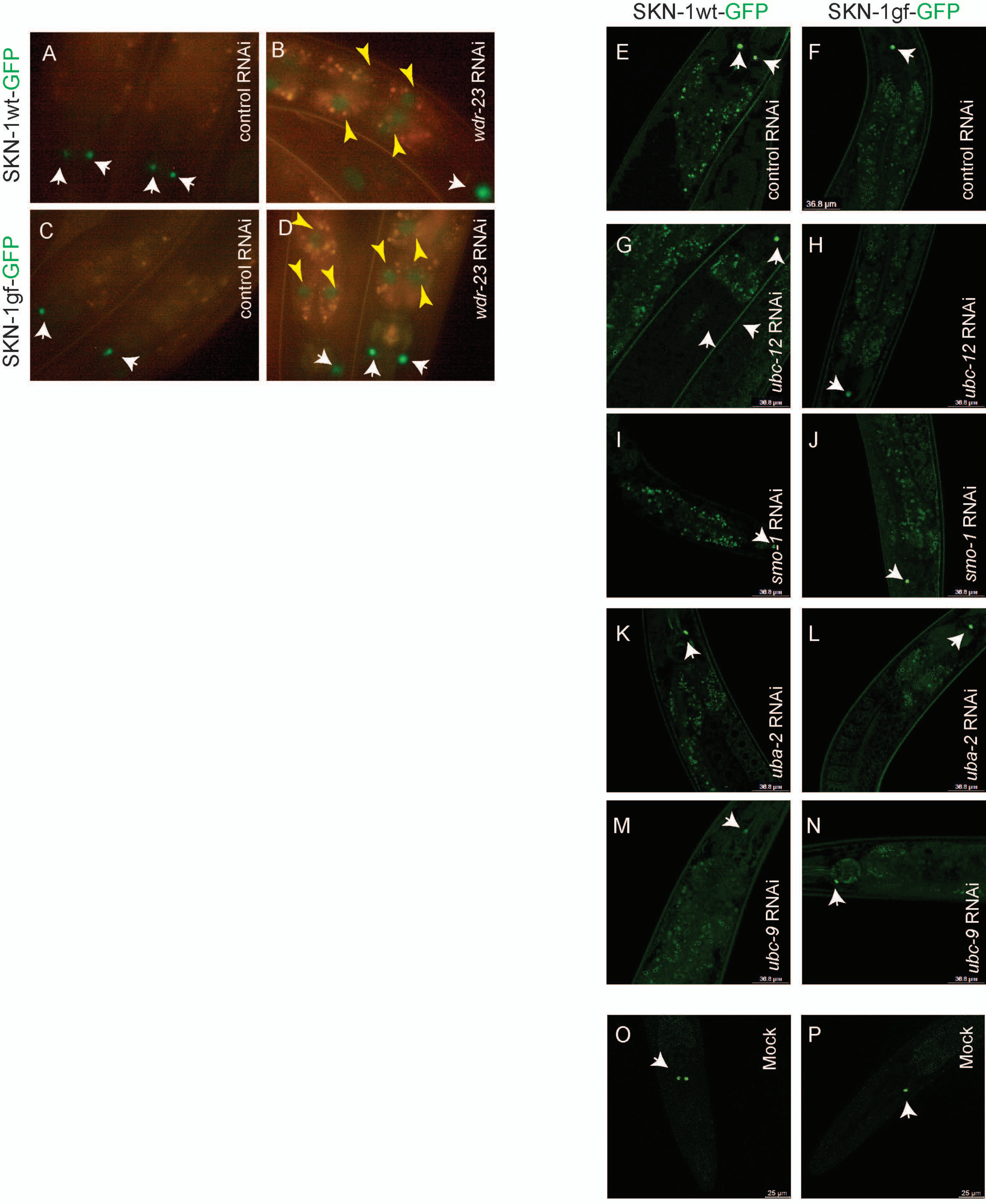


Turner Figure S2

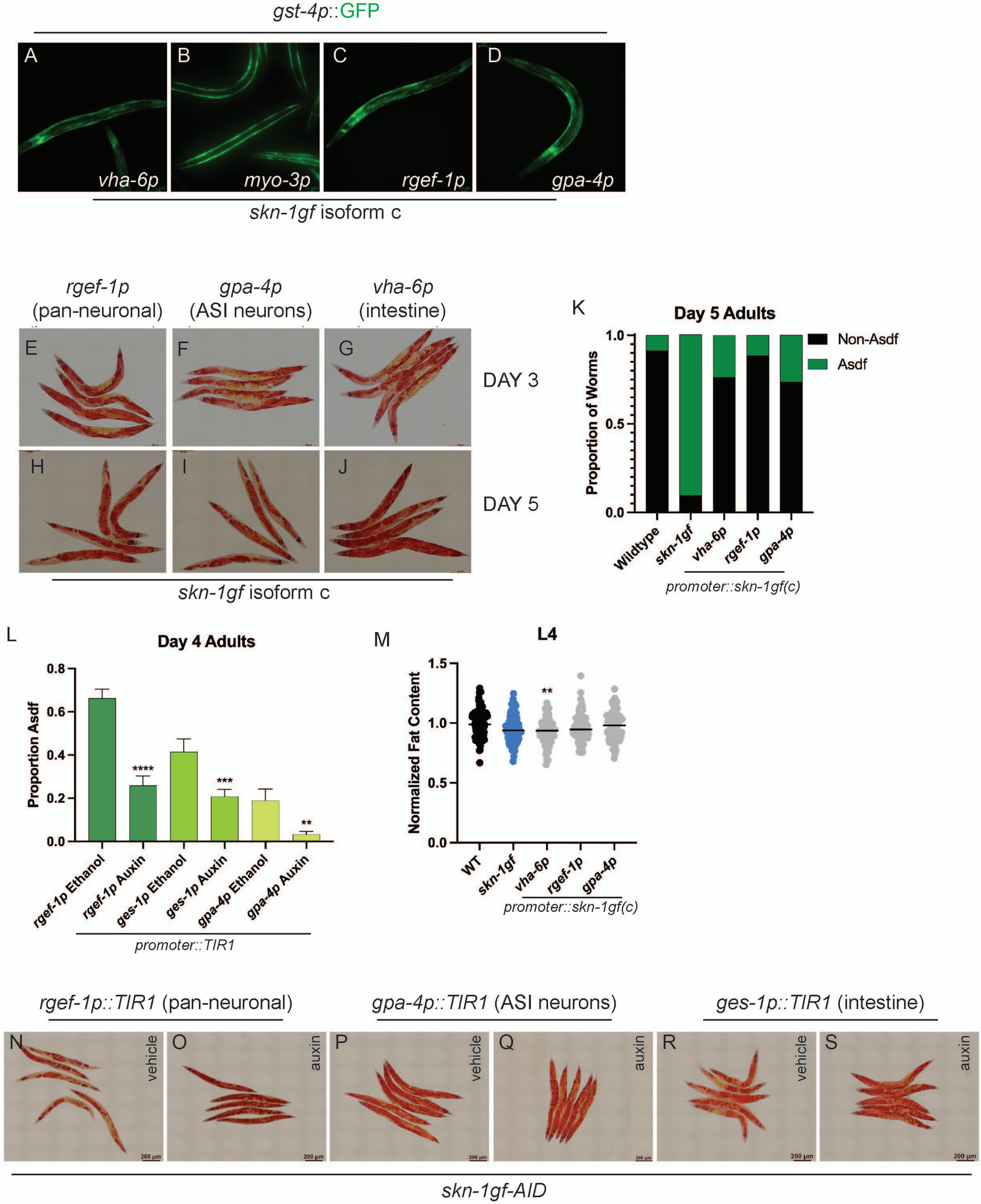

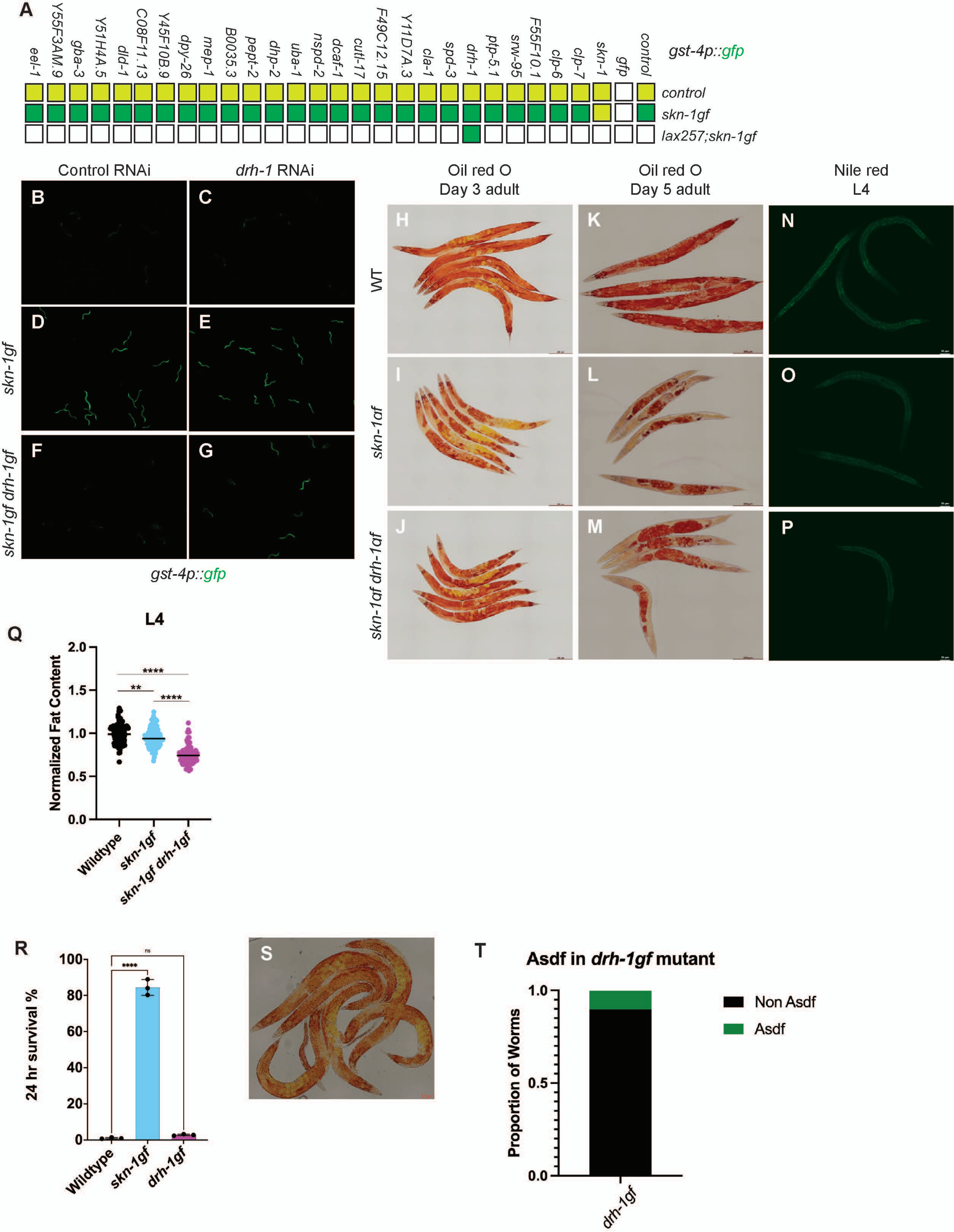

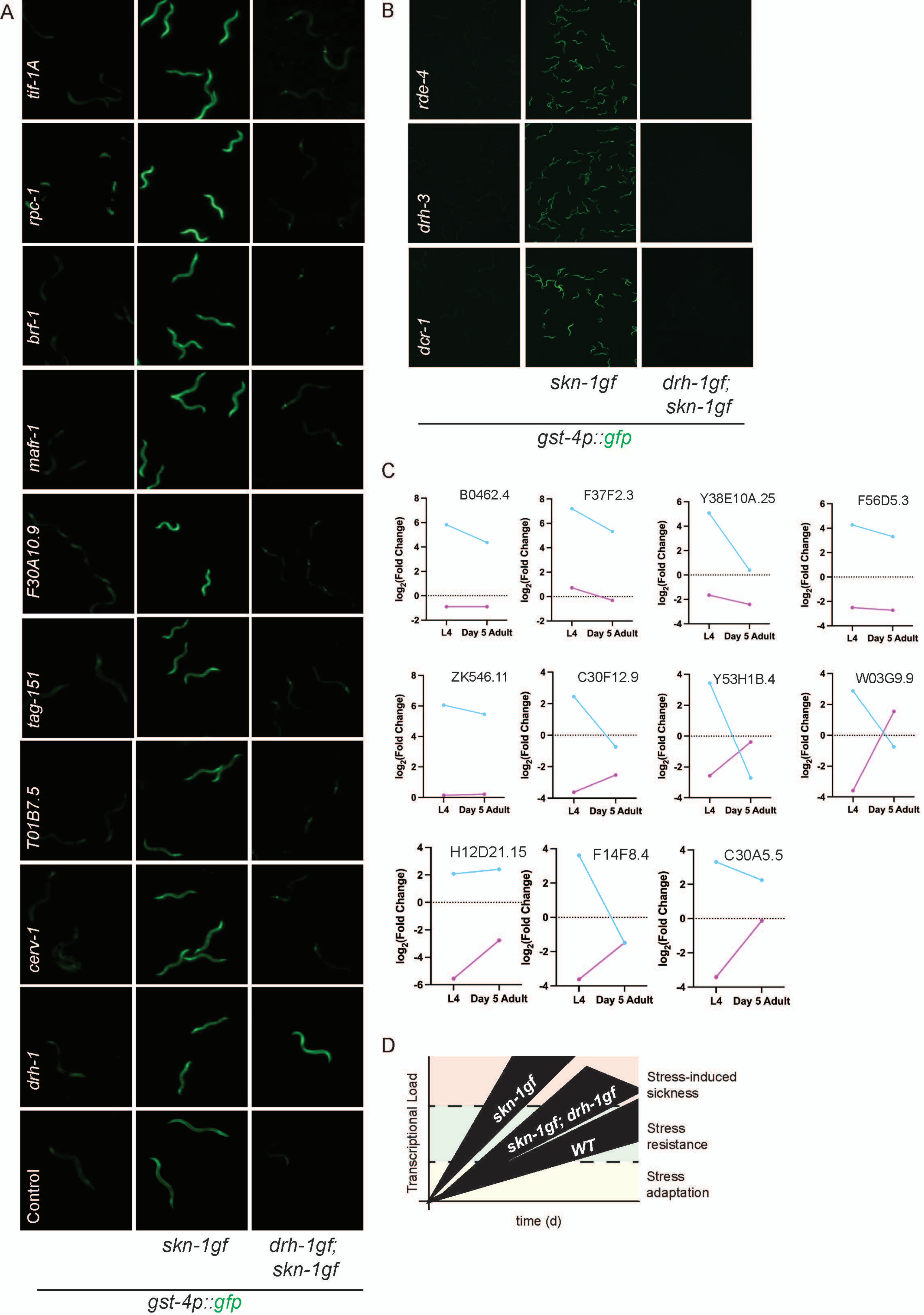
